# Evolutionary profile enhancement improves protein function annotation for remote homologs

**DOI:** 10.64898/2026.03.03.709280

**Authors:** Shitong Dai, Jiaqi Luo, Yunan Luo

**Affiliations:** School of Computational Science and Engineering, Georgia Institute of Technology

**Keywords:** Protein function prediction, Protein language models, Machine learning

## Abstract

Accurate annotation of protein function is essential for understanding biological processes, yet this remains challenging for proteins lacking characterized homologs or belonging to underrepresented functional classes. Although machine learning approaches have become the gold standard for automated function prediction, they often perform poorly on out-of-distribution samples with low sequence identity to training proteins with known annotations. We propose EPERep, an evolutionary input enhancement strategy that leverages the vast space of unannotated protein sequences to improve the prediction of the functions of underrepresented proteins. Our key insight is that, even if a query protein has insufficient similarity to annotated proteins for direct annotation transfer, a wider range of similar unannotated sequences can be identified to facilitate better representation learning. Inspired by profile-based sequence search methods, EPERep incorporates homologous sequences as contextual input to refine the representations of individual proteins from pre-trained protein language models, effectively constructing a pLM-based profile for each query protein. Across four major annotation benchmarks on EC numbers, structural domains, Pfam families, and Gene Ontology predictions, EPERep consistently outperforms strong ML and sequence-alignment baselines. Gains are most pronounced for proteins from rare functional classes, with few or no labeled homologs, and for sequences exhibiting remote homology to the training distribution. These results demonstrate that evolutionary input enhancement provides a principled and scalable strategy for improving protein function prediction, particularly in long-tail and low-identity regimes.

## Background

Proteins underpin virtually all biological processes. With the rapid accumulation of genomic sequences enabled by next-generation sequencing, elucidating the functions of encoded proteins has become a central challenge in modern biology. Computational approaches play a crucial role in automating protein function annotation. Classical sequence search tools such as BLAST^1^, Diamond^2^, MMseqs2^3^, HMMER^4^, and HH-blits^5^ assign annotations to uncharacterized proteins by transferring functions from homologous sequences. However, relying solely on sequence similarity often leads to annotation errors due to the compositional and structural complexity of proteins, such as domain shuffling, fusions, or combination events^6^. Functional diversification driven by duplication and divergence can also yield proteins with distinct functions despite substantial sequence similarity^7^.

To overcome these limitations, machine learning (ML) methods aim to directly capture function similarity beyond sequence similarity by learning complex mappings from sequence to function. These approaches have achieved state-of-the-art performance across diverse annotation tasks^8–10^ and have powered the winning algorithms in recent Critical Assessment of Functional Annotation (CAFA) challenges^11,12^. As a result, major databases such as UniProt and InterPro have begun incorporating ML-based predictions alongside experimentally validated annotations in their official releases.

Despite these advances, ML-based predictors remain unreliable for proteins that lack functionally characterized homologs or belong to underrepresented functional classes. Such cases are challenging because modern ML models implicitly depend on the similarity between the learned representations of the test and training proteins. When a query lies far from all training samples (out-of-distribution scenario), the model cannot confidently assign any function, often performing little better than random guessing. This problem is exacerbated by the intrinsic imbalance of protein function annotation databases such as Gene Ontology (GO), Enzyme Commission (EC) numbers, and Pfam, where a small fraction of well-studied proteins dominate the label space, while the vast majority are sparsely annotated. In our previous work^13^, we showed that standard ML training objectives disproportionately allocate model capacity toward frequent classes, since optimizing these classes most efficiently minimizes the global loss. Consequently, representations of rare or novel proteins tend to be undertrained and less discriminative. Although several recent efforts^14–17^, including ours^13^, have sought to address this imbalance, the accurate annotation of proteins with limited homology or rare functions remains an open challenge.

To address this, we propose an evolutionary input enhancement strategy that enriches the representation of each query protein with evolutionary contextual information, thereby improving function prediction for underrepresented proteins. The UniProt database contains more than 250M protein sequences (Release 2025 03), yet only a small fraction (570K proteins in Swiss-Prot) carry curated functional annotations. Our key observation is that even when a query protein is too dissimilar from annotated proteins for direct label transfer, it often shares a higher similarity with numerous *unannotated* sequences that can nevertheless inform its representation. We thus retrieve homologous sequences for each query and condition the representation learning process on these related sequences. This is achieved by incorporating the retrieved homologs as additional input to refine the sequence representations generated by pre-trained protein language models (pLMs), in contrast to conventional methods that only focus on the query protein.

Our approach is inspired by the evolution of sequence search algorithms. Profile-based methods such as PSI-BLAST^14^, HHblits^5^, and HMMER3^18^ extended the seminal BLAST algorithm by aligning not only single query sequences against other sequences but also incorporating homologous sequences from multiple sequence alignments (MSAs) to build a profile (position-specific scoring matrix) for the query. These profile-based methods have demonstrated excellent sensitivity improvements in remote homology detection. Analogically, here we construct a pLM-based profile for each protein, leveraging homologous sequences to produce richer, context-aware representations that better capture evolutionary and functional relationships.

Building on this design, we comprehensively evaluated EPERep across four major protein function annotation tasks: EC number, structural domain, protein family, and Gene Ontology prediction, where it shows consistent improvements over state-of-the-art sequence- and ML-based baselines. The advantages of EPERep are especially pronounced in challenging but biologically important regimes, including underrepresented functional classes, proteins with few or no annotated homologs, and sequences with low identity to the training distribution. Systematic analyses reveal that EPERep achieves performance gains through two complementary mechanisms: it (i) bridges sequence-identity gaps between distant queries and functionally related training proteins, and (ii) constructs function-coherent evolutionary profiles that highlight subtle sequence features not detectable from single sequences alone. Together, these results position EPERep as a principled and scalable framework for annotating proteins with rare activities or remote homology to known families, and as a general strategy for integrating evolutionary context into modern protein representation learning.

## Results

### Overview of EPERep

We consider the sequence-based protein function annotation problem in this work. Given a protein’s amino acid (AA) sequence as input, the goal is to assign the protein one or more labels from a controlled vocabulary of function annotations. These annotations are typically curated in various databases that describe protein functions in different biological perspectives. Representative examples of protein function annotation system include Gene Ontology^19^ (general biological functions), EC numbers^20^ (enzymatic activities), Pfam^21^ (protein families), Gene3D^22^ (structural domains), as well as other schemes describing protein-protein interactions^23^, sequence domain^24^, conserved residues^25^, and pathways^26^.

EPERep is a sequence-based ML framework for protein function annotation. Given a query protein sequence, EPERep retrieves its homologous sequences from a large protein sequence database (e.g., UniRef30) using MMSeqs2^3^. The *k* most similar homologous sequences, together with the query itself, form an *evolutionary profile* that serves as the model input. The profile is then encoded by a protein language model (pLM)-based neural network to obtain per-sequence embeddings. These embeddings are aggregated into a single contextualized representation of the query protein, which is subsequently passed through a multi-layer perceptron (MLP) classifier to predict function annotations. Unlike conventional methods that rely solely on a single-sequence input, EPERep integrates evolutionary context to enhance latent representations, amplifying subtle but functionally informative sequence features–particularly beneficial for proteins that are remote homologs of annotated sequences.

Formally, let ***s*** = *a*_1_*a*_2_ … *a*_*L*_ denote a protein sequence of length *L*, where *a*_*i*_ ∈ *S* is the amino acid at position *i*, and *S* is the set of 20 canonical AAs. Let ℒ = {1, 2, …, *B*} denote the vocabulary of protein function labels (e.g., GO terms or EC numbers). A protein function annotation database can thus be represented as a collection of protein-function pairs 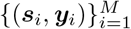 where ***y***_*i*_ is the set of labels associated with sequence ***s***_*i*_. Since proteins may perform multiple functions, ***y***_*i*_ may include more than one label from ℒ.

In general, ML-based protein function annotation can be formulated as a multi-label, multi-class classification problem, where the goal is learning an ML model *f*_*θ*_ : 𝒮 → 2^ℒ^ parameterized by *θ*. Most existing methods are instantiated with deep neural networks^10,16,17,27–29^ that can be decomposed into a feature extractor *h*_*ϕ*_ and a linear classifier *g*_*ω*_, such that *f*_*θ*_ = *g*_*ω*_ ◦ *h*_*ϕ*_. The feature extractor encodes the input sequence to a latent representation ***x*** = *h*_*ϕ*_(***s***) ∈ ℝ^*d*^, and the classifier predicts the function label probabilities ***ŷ*** = *g*_*ω*_(***x***) ∈ ℝ^*B*^. Unlike alignment-based methods that transfer annotations based on raw sequence similarity, these ML models predict protein functions by exploiting function similarity captured by representation ***x***. However, when the query sequence exhibits very low identity to all training proteins, the learned representation may be poorly grounded, resulting in unreliable predictions.

EPERep mitigates this limitation by augmenting the input sequence with its evolutionary contexts of homologous sequences to amplify the low signals of sequence similarity, facilitating information-rich representation learning and guiding the function prediction. Given an given protein ***s***, we first uses MMSeqs2^3^ to retrieve its *k* most similar sequences ℛ (***s***) = {***r***_1_, …, ***r***_*k*_} from an external sequence database (UniRef30; Methods) and define its evolutionary profile as 𝒫 (***s***) = {***s***} ∪ ℛ (***s***). Each sequence in 𝒫 (***s***) is first embedded using ESM-2^30^, a large-scale pre-trained pLM that captures evolutionary and structural constraints from millions of protein sequences. The resulting embeddings are further refined using ProteinCLIP^31^, a bimodal pLM jointly trained on protein sequences and natural-language descriptions to align structural and functional semantics. In this work, we retrained a ProteinCLIP model on its original training set, excluding sequences in our test data to avoid data leakage through natural-language descriptions. An aggregation module based on the multi-head attention architecture^32^ to integrate information from all profile members to produce a single contextual representation of the query protein, conditioned on its homologs (Methods). This representation is then passed to fully connected layers to predict functional labels. During training, only the parameters of the aggregation and prediction modules are optimized using cross-entropy loss, while the pre-trained ESM-2 and ProteinCLIP remain frozen, which allows parameter-efficient training across annotation tasks of varying scales.

The core innovation of EPERep lies in the representation learning stage, where the feature extractor learns from both the query sequence ***s*** and its homologous context ℛ(***s***), i.e., ***x*** = *h*_*ϕ*_({***s***}∪ℛ(***s***)). Here, *h*_*ϕ*_ is a pLM that encodes the query ***s*** and homologous contexts ℛ(***s***)} into embeddings, followed by an aggregation module that summarizes these embeddings into a unified embedding. Akin to how profile-based search algorithms (e.g., PSI-BLAST, HMMER) extend pairwise comparison methods (e.g., BLAST) by constructing multiple-sequence profiles to improve sensitivity in remote homology detection, EPERep generalizes existing sequence-based ML models through evolutionary input enhancement to boost function annotation accuracy for novel proteins. By leveraging pLMs to construct sequence profiles that capture higher-order dependencies and long-range constraints, EPERep generalizes traditional MSA- or HMM-based profiles that model only site-specific preferences or first-order sequence dependencies. Notably, incorporating evolutionarily related sequences is central to the success of structure prediction models like AlphaFold^33^. Our work differs in focusing on learning sequence-level representations optimized for function annotation, whereas AlphaFold learns residue-level geometric representations for structure prediction. EPERep is also closely related to the emerging trend in protein or genomic language models that leverage MSAs to enrich sequence representation learning^34–45^, while their utility for enhancing function annotation for remote homology protein sequences remained underexploited.

### Accurate predictions for diverse protein function annotation tasks

To evaluate the performance of EPERep, we collected the benchmark datasets introduced in MSRep^13^ covering four function annotation tasks: EC numbers, Gene3D codes, Pfam families, and GO terms (Fig. 2a). For each task, protein sequences and function annotations were sourced from the Swiss-Prot database^46^. We used a time-based train-test split with May 25, 2022 as the cutoff and removed all test proteins sharing more than 50% sequence identity with any training protein, resulting in challenging test sets with both temporal and sequence identity controls. Model performance was evaluated using AUPR and Fmax, following the established CAFA protocols^11^.

**Figure 1:**
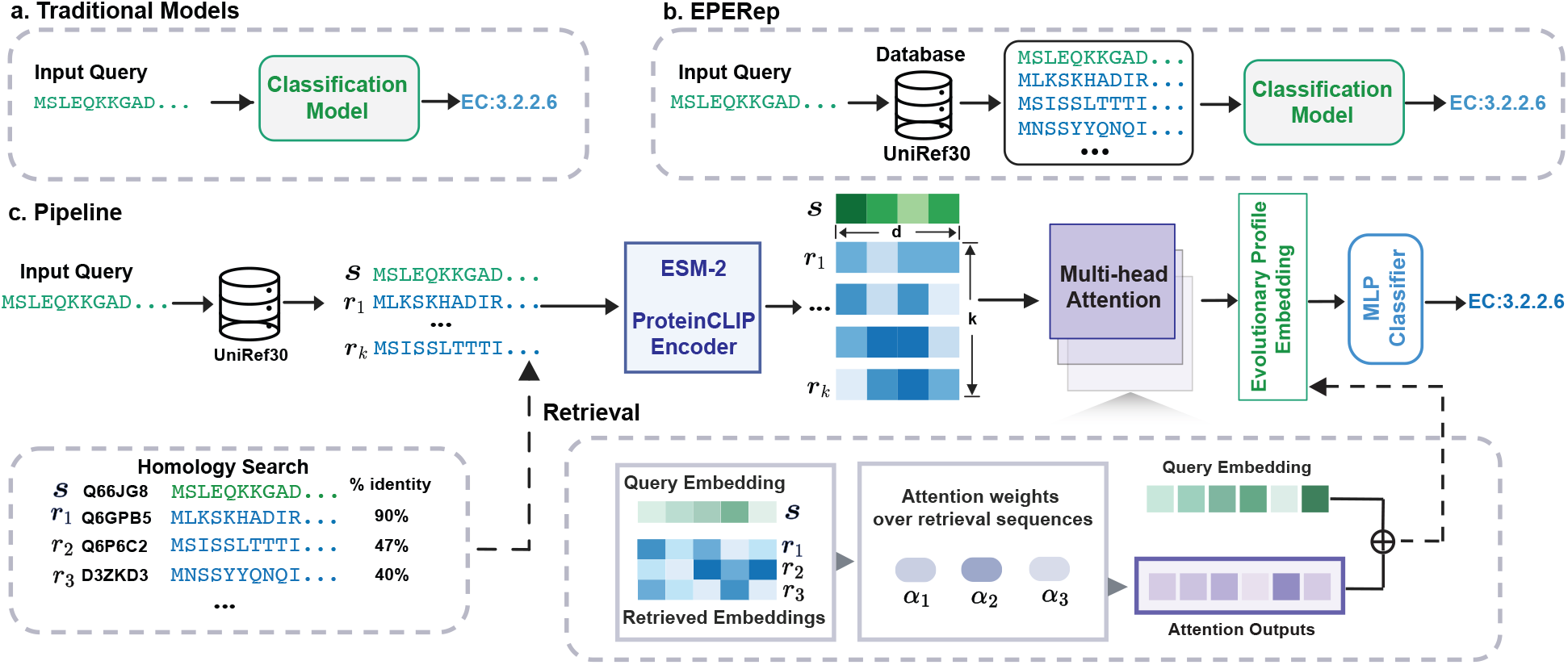
Overview of EPERep. **(a) (b)** Conceptual comparison between traditional single-sequence models and EPERep. **(a)** Architecture of traditional ML-based protein function prediction methods, where the query protein is treated as an isolated input. **(b)** New paradigm in EPERep, where the query protein is enhanced with homologous sequences from a large, unannotated database like UniRef30. **(c)** End-to-end pipeline of EPERep. The query and retrieved sequences are encoded with a protein language model, ESM-2, followed by ProteinCLIP, which further aligns the representation with natural language descriptions of proteins. The attention weights of the representations of the retrieved sequences against the query sequence’s representation are computed and used to aggregate the retrieval information. The aggregation outputs are added to the query embedding to form a contextualized representation, followed by a lightweight MLP classifier to produce the final predictions. The protein language encoders are kept frozen during training, while the attention module and MLP classifier are optimized. MLP: multi-layer perceptron.

**Figure 2:**
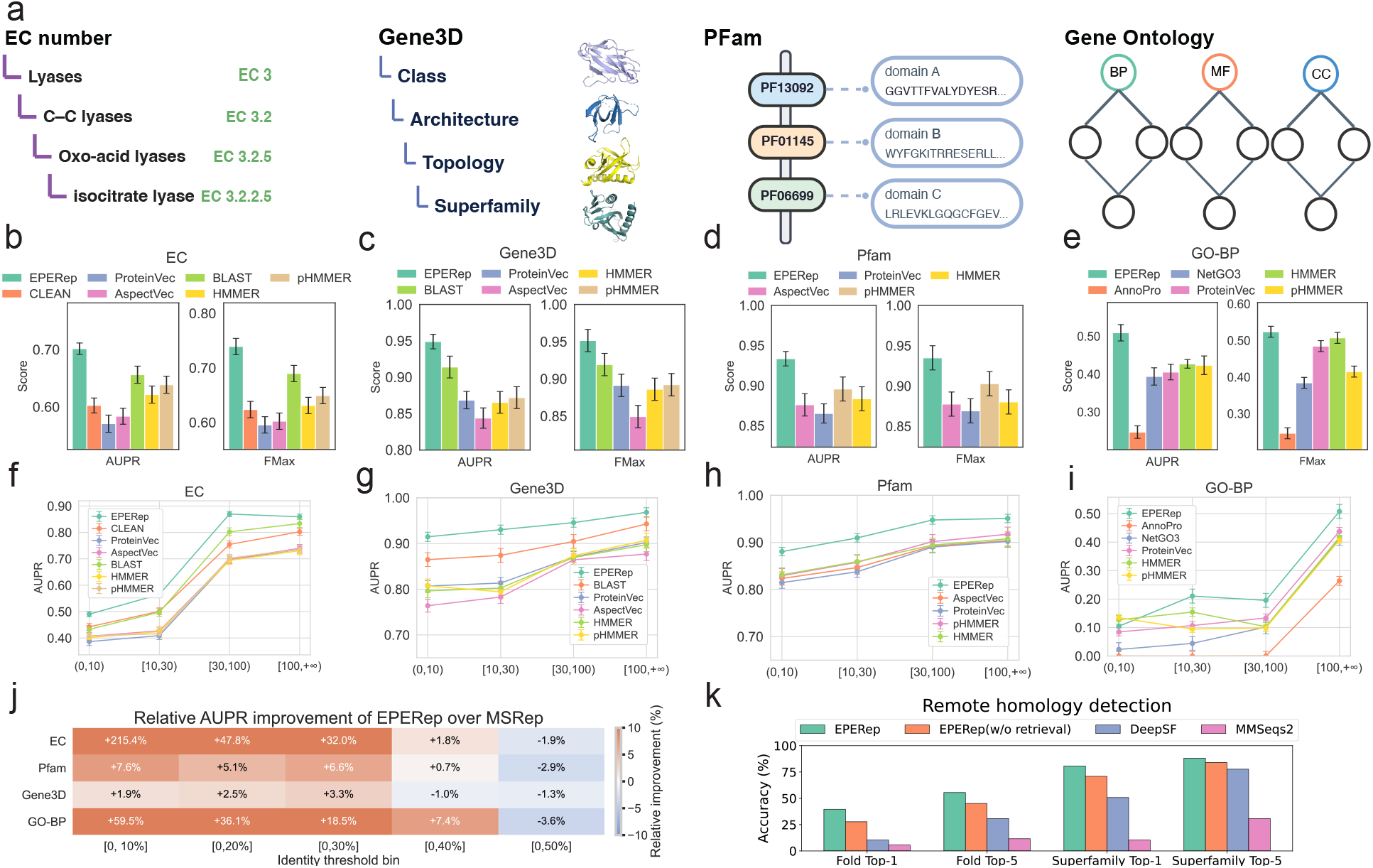
Performance of EPERep across four function annotation tasks. **(a)** Schematic illustrations of the four protein function annotation tasks: EC numbers, Gene3D structural domains, Pfam families, and Gene Ontology (GO). **(b-e)** AUPR and Fmax scores of EPERep and baseline methods on the four annotation tasks, respectively. BP: biological process; MF: molecular function; CC: cellular component. **(f-i)** Stratified AUPR scores for EPERep and baselines across subsets of function labels grouped by their frequencies in the training set (number of proteins associated with the label): [0, 10), [10, 30), [30, 100), [100, +∞). Bar plots in (b–e) and line plots in (f–i) show mean *±* s.d. over 10 rounds of 90% bootstrap resampling of the test sets. **(j)** Dot plot of MSRep vs. EPERep by sequence-identity bin. For each test protein, we search the training set and take the maximum sequence identity (max identity) to any training protein (i.e., the best-hit identity). The x-axis shows cumulative bins of this value (e.g., [0, 0.3] includes all test proteins whose best hit in the training set has identity ≤ 0.30). Red indicates where EPERep outperforms MSRep, and blue bins indicate the opposite; the lighter shade shows the other model’s score. **(k)** Remote homology detection performance on the DeepSF benchmark. Top-1 and Top-5 accuracies at both the fold and superfamily level are shown for EPERep, baseline models, and ablated variants.

#### EC numbers

EC number is a four-level hierarchical system that categorizes enzymatic reactions with increasing specificity. EPERep directly predicts the finest-grained, four-digit EC numbers. Compared with recently developed deep learning predictors (CLEAN^10^, Protein-Vec^17^, Aspect-Vec^17^), traditional sequence-to-sequence search tools (BLAST^1^ and pHMMER^47^), and profile-to-sequence search methods (HMMER^4^), EPERep achieves the best overall performance in AUPR and Fmax (Supplementary Note A.1). While all deep learning baselines are substantially outperformed by BLAST, EPERep shows 2.7% higher AUPR and 2.9% higher Fmax over BLAST (Fig. 2b), suggesting the effectiveness of input augmentation with evolutionary contexts.

#### Gene3D

Gene3D codes provide structural domain annotations for protein sequences, with a four-level hierarchy from CATH^48^: Class, Architecture, Topology, and Homologous superfamily. A protein may be annotated with more than one Gene3D label since it can include multiple structural domains. We evaluated EPERep on the most specific Homologous superfamily level. Compared with Protein-Vec, Aspect-Vec, and BLAST, the latter of which is highly competitive on this task, EPERep achieves further improvements in both AUPR and Fmax (Fig. 2c). These results demonstrate that EPERep not only matches but improves upon sequence similarity–based methods by leveraging homologous context.

#### Pfam families

Pfam organizes protein sequences into curated families defined by MSAs and profile HMMs. Compared with the EC number and Gene3D tasks (each with ~5,000 classes), Pfam represents a substantially larger and more challenging label space of over 14,000 families. Nevertheless, EPERep consistently outperformed ML models (Protein-Vec, Aspect-Vec) and sequence search tools (HMMER and pHMMER), improving AUPR and Fmax by 5.5% and 6.9%, respectively (Fig. 2d). These results indicate that evolutionary input enhancement scales effectively to high-cardinality annotation schemes.

#### Gene Ontology

The Gene Ontology (GO) organizes biological knowledge into a directed acyclic graph of functional terms, creating a substantially more complex structure than the hierarchical trees of EC or Gene3D annotations. This complex ontology structure makes the prediction of GO one of the fundamental challenges in bioinformatics^11^. Following the common practices^11^, we trained three separate EPERep models for the three sub-ontologies of GO: biological process (BP), molecular function (MF), and cellular component (CC). Across the three categories, EPERep outperforms leading ML methods (AnnoPro^49^, NetGO3.0^50^, and Protein-Vec) and alignment-based methods (Figs. 2e and S1a-d).

### Enhanced annotations for understudied functions and remote homologs

We found that the gains of EPERep over existing approaches are most pronounced in two challenging but biologically important scenarios: (1) predicting functions that are severely underrepresented in current databases, and 2) annotating proteins that are remote homologs of functionally characterized sequences.

Most protein function annotation datasets exhibit extreme class imbalance, where a small portion of well-studied function labels are associated with most proteins, whereas the majority of labels appear in only a handful of proteins (e.g., *<* 10). To evaluate whether EPERep maintains predictive accuracy across this long-tail distribution, we stratified test proteins by the frequency of their associated function labels in the training set into four bins: [0, 10), [10, 30), [30, 100), [100, +∞). Across four annotation tasks and sixteen total bins, EPERep achieved the highest AUPR in 15 of them (Fig. 2f-i). As expected, all methods experienced reduced accuracy on low-frequency labels; however, EPERep exhibits the smallest performance drop. These results highlight that incorporating evolutionary context via retrieval augmentation mitigates class-imbalance effects by exposing the model to informative homologous sequences even when annotated training examples are scarce.

Our previous work, MSRep^13^, also sought to address the above long-tail problem by learning maximally spanning representations that enhance discrimination for classifying understudied functions. While MSRep improves annotation for low-frequency labels, it relies solely on single-sequence inputs. By contrast, EPERep additionally leverages evolutionarily related sequences to construct enriched representations. We hypothesized that this design would also improve predictions for proteins lacking close homologs in the annotated training set. To test this, we binned test proteins by their maximum sequence identity to any training sequence using cumulative thresholds [0, *x*%] with *x* = 10, 20, …, 50. EPERep consistently outperforms MSRep for low-identity proteins (≤ 30% identity), with higher performance gains observed when query sequences are more distant from training sequences (Fig. 2j). These results support the intuition that evolutionary profiles help recover functional signal for difficult low-identity queries where single-sequence models struggle. For higher identity bins (40–50%), EPERep and MSRep achieve comparable performance, likely because single-sequence representations already contain sufficient evolutionary signals to inform function prediction, leaving less room for retrieval-based enhancement. Together, these findings indicate that EPERep is complementary to MSRep, particularly in scenarios where annotated homologs are sparse or absent.

Motivated by these observations, we further assessed EPERep’s generalizability on a classical remote homology detection benchmark. Remote homology detection evaluates whether a model can identify relationships between proteins that share the same structural fold despite low sequence similarity. We used the SCOP 1.75 dataset^51^, which organizes proteins into a Class/Fold/Superfamily/Family hierarchy. Following DeepSF^52^, we trained EPERep to predict a sequence’s Fold-level label under two strict data splits where any two training and test sequences shared at most Fold- or Superfamily-level labels, while sharing no labels beyond the indicated level. Across both splits, EPERep achieved substantial improvements over DeepSF^52^ and MMSeqs2 (Fig. 2k). Top-1 accuracy increased by 29.3% and Top-5 accuracy by 24.6%. Ablation experiments further revealed that removing the retrieval augmentation module reduces accuracy by 12–14%, underscoring that evolutionary profiles provide informative cues even in the extremely low-identity regime characteristic of remote homologs. These results demonstrate that retrieval-enhanced representations learned by EPERep can reliably capture deep evolutionary relationships beyond those accessible from single sequences.

### Retrieval augmentation provides evolutionary context beyond the training set

Having evaluated EPERep’s performance, we aimed to understand the source of its performance gains by analyzing its behavior on the EC number prediction task. We grouped into four categories based on whether EPERep and Aspect-Vec (an ML baseline that does not use retrieval) correctly predicted their labels: TT, FT, TF, and FF (e.g., TF includes proteins correctly predicted only by EPERep; Fig. 3a). For each protein, we computed two sequence identity measures: (i) the mean identity to its top-*k*=10 nearest neighbors retrieved from UniRef30, and (ii) the mean identity to its top-*k* nearest neighbors in the training set. Since traditional single-sequence-input models can only exploit information from the training set, (ii) represents the best sequence similarity signal accessible to those baselines.

**Figure 3:**
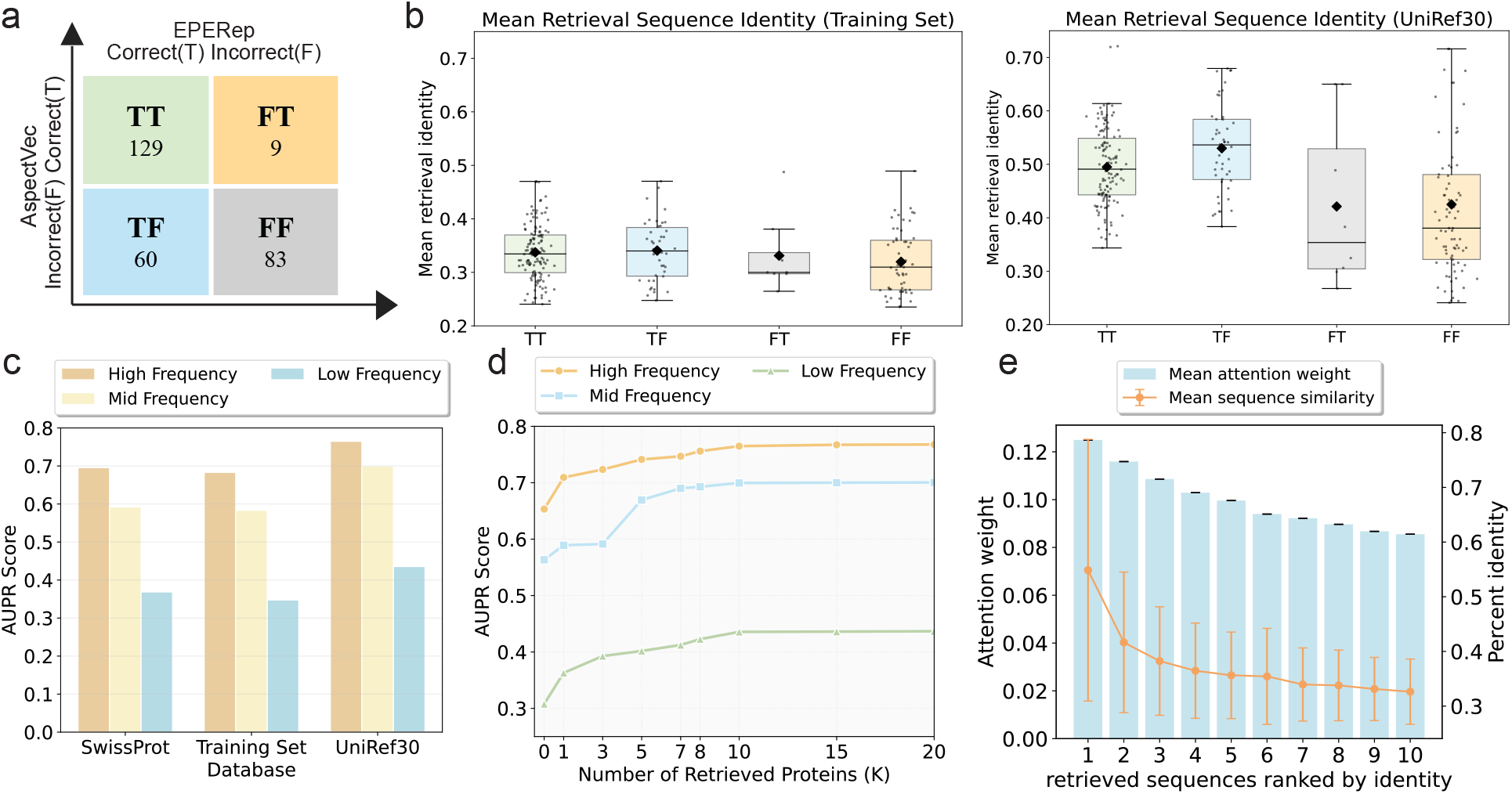
Effects of retrieval augmentation in EPERep. Results shown are for the NEW-2022 test set of EC number predictions. **(a)** Prediction outcomes of EPERep and Aspect-Vec. Each cell corresponds to one of four groups: TT (both correct), TF (EPERep correct only), FT (Aspect-Vec correct only), and FF (both incorrect). Counts indicate the number of test proteins in each group. **(b)** Mean sequence identity between each test protein and its top-*k* neighbors retrieved either from the training set or from UniRef30. The x-axis follows the four prediction groups in (a). **(c)** AUPR performance of EPERep when using retrieval databases of increasing scale (Swiss-Prot, EC training set, and UniRef30), stratified by label frequency in the training data: Low ([0, 30)), Mid ([30, 100)), and High ([100, +∞)). **(d)** Effect of the number of retrieved sequences *k* on EC prediction performance, evaluated separately within the low-, mid-, and high-frequency label groups. **(e)** Relationship between attention weights assigned to retrieved neighbors and their sequence identity to the query. Attention scores are averaged across all attention heads in the aggregation module.

We found that for all four groups, the mean identities to the top-*k* training-set neighbors fall within the range of 30%-35% (Fig. 3b). In contrast, retrieval from UniRef30 yields substantially more similar sequences, especially for proteins in the TF group, where EPERep succeeds but Aspect-Vec fails. These proteins exhibit retrieved neighbors with *>*50% identity on average, representing a dramatic improvement over the ~35% identities available within the training set. Thus, the evolutionary profile augmentation enables EPERep to incorporate more informative homologs that are absent from the annotated training data. Importantly, even though retrieved sequences may share EC numbers with the query, only their *sequences* are used, and the model never accesses their functional labels (if any), avoiding data leakage and ensuring that performance improvements arise purely from richer sequence context.

To further quantify how retrieval database size influences performance, we trained EPERep using three search databases of increasing scale: the EC training set (200k sequences), Swiss-Prot (570k), and UniRef30 (200M) and compared their stratified prediction performance for EC numbers with High/Mid/Low frequencies (defined in Fig. 3 caption) in the training data. The result reveals that performance improves as the retrieval database grows (Fig. 3c). The gains are largest for low-frequency EC labels, moderate for mid-frequency labels, and smallest for high-frequency labels. This trend reinforces that evolutionary input enhancement is most beneficial when conventional models underperform, such as when annotated homologs are scarce.

We also examined how performance varies with the number of retrieved homologs *k*. On the EC task, accuracy increases steadily from *k* = 0 to *k* = 10 (Fig. 3d). The marked improvement from *k* = 0 to *k* = 1 highlights the benefit of even a single retrieved sequence over query-only representations. Performance plateaus beyond *k* = 10, indicating that this choice offers a favorable balance between information richness and computational efficiency. Furthermore, attention weights learned by the aggregation module strongly correlate with the sequence identity between each retrieved neighbor and the query (Fig. 3e), confirming that EPERep adaptively prioritizes the most informative homologs.

Together, these analyses demonstrate that EPERep’s improvements stem from both the *scale* and *quality* of retrieval. Access to large databases such as UniRef30 substantially increases the likelihood of finding highly similar or functionally relevant sequences that are absent from curated training sets. Combined with a learnable attention mechanism that selectively integrates the most informative neighbors, EPERep effectively constructs enriched evolutionary profiles that enhance representation learning and function prediction accuracy.

### Two complementary mechanisms driving evolutionary profile enhancement

We identified two complementary mechanisms by which the retrieval augmentation module in EPERep improves function prediction: *sequence-level bridging* and *profile-level enrichment*. Together, these mechanisms enable the model to construct a richer, evolutionarily grounded context for each query protein, allowing more accurate predictions even for sequences that are highly divergent from the labeled training distribution.

#### Sequence-level bridging

ML models rely fundamentally on similarities between unseen queries and training examples. Under the strict sequence-identity constraint used in this study (*<*50% identity between any training–test pair), many queries become out-of-distribution instances where direct similarity to annotated proteins is too weak to be informative. EPERep addresses this problem by creating an evolutionary profile whose members often exhibit substantially higher similarity to the query than any annotated sequence in the training set.

Across the EC test set, Fig. 4b shows that retrieved neighbors from UniRef30 have markedly higher sequence identity and embedding similarity to functionally matched training proteins than to unmatched ones, indicating that retrieval naturally prioritizes proteins enriched in relevant functional signals.

**Figure 4:**
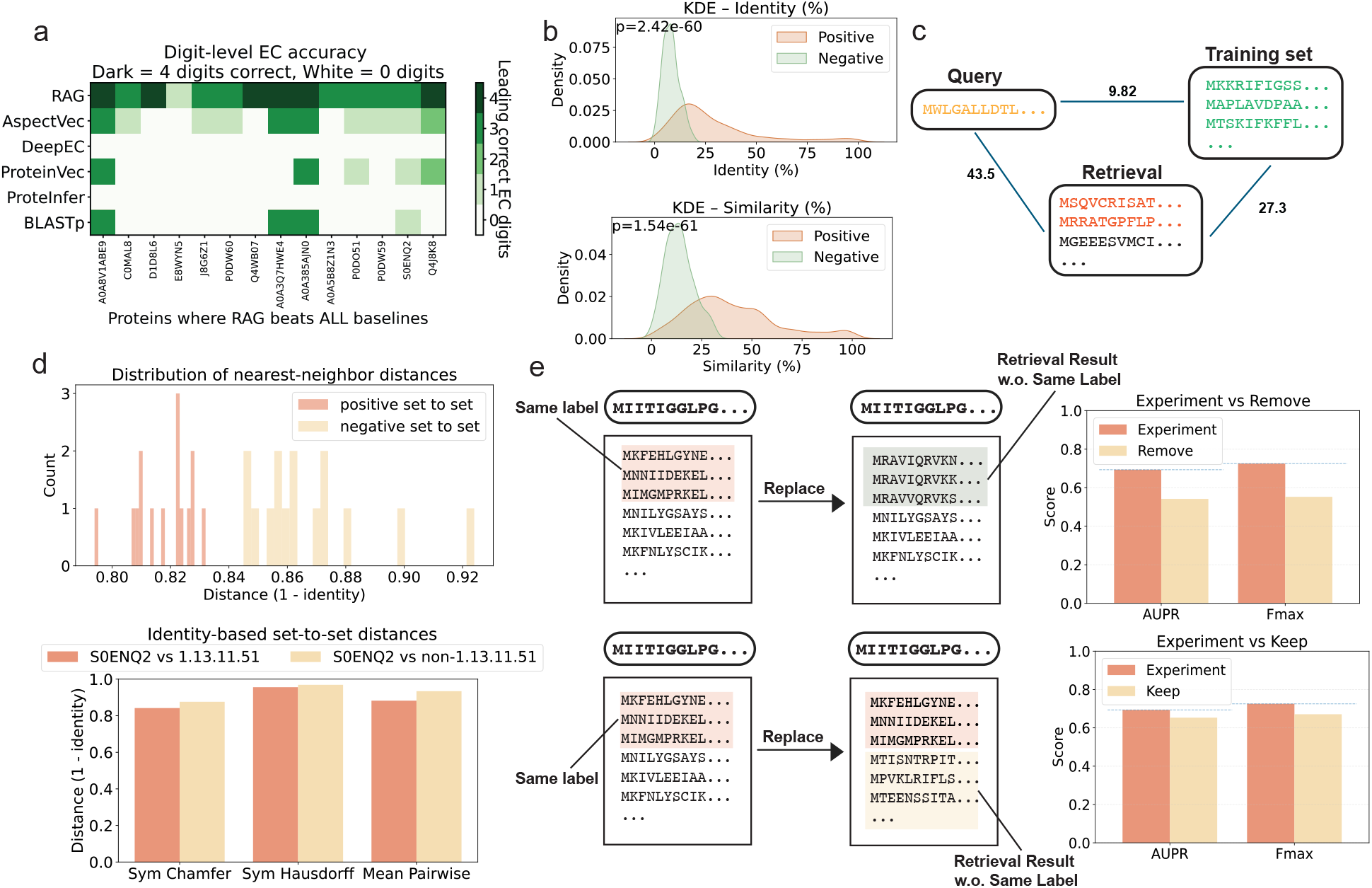
Case studies and ablation analyses of evolutionary profile retrieval. **(a)** Test proteins on which EPERep outperforms all baselines in EC digit-level accuracy. Darker shades indicate a greater number of leading EC digits correctly predicted (4 = perfect, 0 = none). **(b)** Kernel density estimates (KDEs) of global-alignment sequence identity and embedding similarity between retrieved neighbors and training proteins. “Positives” denote training proteins sharing the query’s EC label; “negatives” do not. **(c)** Average pairwise sequence identity among three protein groups for query `J8G6Z1`: (1) query vs. same-label training proteins, (2) query vs. retrieved neighbors, and (3) retrieved neighbors vs. same-label training proteins. **(d)** Set-to-set distances for query `S0ENQ2`. Positive distances measure similarity between the retrieved set and the set of training proteins sharing the same EC label; negative distances compare to randomly sampled training proteins with different labels. Distances are computed using symmetric Chamfer, symmetric Hausdorff, and mean pairwise metrics. **(e)** Label-consistency ablations at fixed *k*. In the label-excluded setting, all same-label neighbors are removed and replaced with the next closest non–same-label hits. In the label-retained setting, same-label neighbors are preserved while remaining retrieved proteins are replaced with randomly sampled neighbors. Protein-centric AUPR and Fmax scores are shown for both ablations.

Two illustrative cases further clarify this mechanism. The protein `J8G6Z1` shares only 9.8% mean identity with training proteins of the same EC number, leading baseline models to entirely fail. EPERep correctly predicts two EC digits because the retrieved neighbors narrow this gap: they share 43.5% identity with the query and 27.3% identity with the correct-label training proteins (Fig. 4c). This effectively creates a “bridge” that carries functional information from distant training proteins to the query. The protein `S0ENQ2` presents a different pattern: none of its retrieved neighbors share its EC label. Yet EPERep still outperforms all baselines because the retrieved sequences remain substantially closer (in both sequence and embedding space) to the positive training group than to randomly chosen negatives (Fig. 4d). In other words, even imperfectly matched neighbors can form an informative intermediate profile that improves the representation of the query.

These analyses show that retrieval enables EPERep to detect weak but functionally meaningful evolutionary signals by bridging gaps between distant queries and annotated proteins.

#### Profile-level enrichment

Retrieval augmentation also improves prediction through a second mechanism: enriching the representation of the query using collective patterns from its evolutionary profile. This parallels how profile-based search tools such as PSI-BLAST and HHblits outperform pairwise sequence alignment by leveraging multi-sequence profiles that emphasize conserved and functionally important features.

Although not all retrieved sequences share the true EC label, the presence of functionally relevant neighbors is essential. To evaluate their role, we performed two controlled ablations at fixed *k* (Fig. 4e). In the *label-excluded* setting, all neighbors of the same-label were removed and replaced with unrelated sequences of similar identity. This caused sharp performance declines (AUPR −0.15, Fmax −0.17), demonstrating that functionally aligned neighbors provide critical discriminative signals. In contrast, in the *label-retained* setting where same-label neighbors were preserved and all others replaced with random proteins, performance decreased only slightly (AUPR −0.04, Fmax −0.05).

These results show that once the retrieval profile includes a small number of functionally consistent proteins, the remaining neighbors primarily contribute robustness rather than serving as the primary discriminative source. The evolutionary profile therefore acts not just as a bridge but as a functionally coherent representation of the query protein that captures collective sequence features beyond what a single sequence can express.

Together, these results indicate that evolutionary profile enhancement improves protein function prediction through two synergistic mechanisms. First, sequence-level bridging enables information flow between evolutionarily distant proteins by exposing the model to more similar homologs absent from the training set. Second, profile-level enrichment constructs a richer, functionally informed representation that enhances the detectability of subtle sequence signals. These complementary effects explain the consistent performance gains of EPERep across annotation tasks, including particularly challenging low-identity and long-tail regimes.

## Discussion

The rapid expansion of genome sequencing has generated an unprecedented volume of protein sequences, yet functional annotation remains a central bottleneck in translating sequencing data into biological insight. Although large-scale protein language models (pLMs) have substantially improved sequence representation learning, their predictive performance ultimately depends on the evolutionary and functional signals captured in curated training datasets, which represent only a small fraction of the global protein space. In practice, many newly sequenced proteins–particularly those from non-model organisms, environmental samples, or poorly characterized clades–lack close homologs with experimentally validated functional annotations. This creates an “evolutionary context gap”: a query protein may reside in a dense neighborhood of homologous proteins in the sequence space, yet remain distant from any labeled sequences available during supervised training. As a result, even capable pLM-based ML models struggle in long-tail and low-identity regimes that dominate real-world genome annotation tasks^10,49^.

EPERep addresses this gap by augmenting each query protein with its homologous sequences. This design is conceptually analogous to the transition from sequence-to-sequence alignment tools such as BLAST^1^ to profile-based alignment methods such as HMMER^4^, where the incorporation of MSAs substantially improves the sensitivity of detecting remote homologies. Here, EPERep extends this principle into pLM-based protein function annotation. Unlike classical MSA profiles that model position-specific substitution patterns, the pLM-integrated profile captures higher-order sequence dependencies encoded by pretrained models, enabling refinement of query representations beyond what can be achieved from single-sequence inference alone.

Our results demonstrate that evolutionary profile enhancement yields great benefits in biologically important but technically challenging settings. Across EC number, Gene3D, Pfam, and Gene Ontology benchmarks, performance gains are most pronounced for low-frequency functional classes and for proteins with limited similarity to annotated training sequences. Such cases are common in orphan genes, lineage-specific expansions, and proteins from newly sequenced microbial or environmental genomes^53,54^, where experimental characterization remains sparse. By leveraging unlabeled evolutionary neighborhoods, EPERep mitigates the intrinsic class imbalance of existing annotation databases, in which a small set of well-studied functions dominate the labeled data.

Within the broader landscape of pLMs and genomic language models (gLMs), EPERep and a growing body of MSA-based approaches^34,43–45^ illustrate a generalizable design principle for integrating foundation models with large-scale biological sequence repositories. While many current applications of pLMs or gLMs rely on single-sequence inference, optionally followed by task-specific fine-tuning^10,13^, retrieval augmentation introduces dynamic, input-dependent context at inference time, effectively coupling pretrained models to the vast sequence space. This paradigm resembles retrieval-augmented generation in natural language processing^55,56^, where the input prompt is augmented with relevant information retrieved from external sources. As such, EPERep demonstrates that pLMs or gLMs need not operate in isolation; instead, they can be systematically combined with the vast, continually expanding corpus of unannotated sequences.

Despite these advantages, several limitations of EPERep warrant consideration. First, retrieval-based methods depend on the composition, coverage, and quality of the underlying sequence database. When homologous neighborhoods are sparse or highly divergent, contextualization may provide limited additional evolutionary signals. Second, retrieval parameters, such as e-value thresholds, clustering levels, and database choice, may influence the balance between sensitivity and specificity. Although our analyses indicate robustness across settings and diminishing returns beyond a modest number of neighbors, systematic evaluation under identity-controlled or taxonomically restricted retrieval scenarios would further clarify the method’s performance. Third, to ensure scalability and parameter efficiency, we freeze the pretrained pLM encoders. While this design facilitates application across diverse annotation tasks, future work integrating more expressive MSA-based encoders or jointly fine-tuned architectures^34,42,44^ may further enhance performance, particularly for complex ontologies such as Gene Ontology.

## Methods

### Benchmarking datasets

We evaluated our method on the same benchmark datasets used in MSRep^13^, encompassing four protein function annotation tasks: EC numbers, Gene3D codes, Pfam families, and Gene Ontology (GO) terms. These datasets were derived from the expert-curated database Swiss-Prot^46^. We employed a time-based split with a cutoff date of May 25, 2022, simulating realistic evaluations on unseen proteins. To further reduce redundancy, the test sets were filtered using MMSeqs2^3^ to exclude any protein sharing ≥ 50% sequence identity with proteins in the training set.

### Retrieval augmentation

To enrich each query protein with information from homologous sequences, we developed a retrieval augmentation module for EPERep. Specifically, we leveraged MMSeqs2^3^, a fast and sensitive sequence alignment tool, as the search engine for homolog retrieval. For the lookup database, we selected UniRef30^57^, which strikes a balance between scale and redundancy by clustering the complete UniProtKB into representative sequences at 30% identity. This design retains broad evolutionary diversity while eliminating near-duplicate proteins. Compared with smaller curated resources such as Swiss-Prot, UniRef30 offers substantially greater sequence coverage, thereby increasing the likelihood of retrieving homologs that provide additional functional context for the query protein.

For each input protein ***s***, EPERep utilizes MMSeqs2 to retrieve the top-k distinct sequences ***r***_1_, …, ***r***_*k*_ with the highest sequence identities from UniRef30 as the augmented neighbors. We applied an e-value cutoff of 10^−5^ to ensure that only statistically significant sequence alignments were retained. This stringent threshold reduces spurious matches while preserving true homologs, thereby improving the reliability of the retrieved neighbors used for augmentation. The retrieval augmentation process can be formulated as follows, where ℛ denotes the retrieval module:

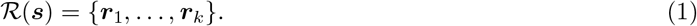

Retrieved entries may or may not carry functional annotations. In all cases, the model has access only to their amino-acid sequences and is agnostic to any labels. This design treats retrieval as unlabeled contextual evidence, reducing spurious matches via the stringent cutoff while preserving true homologs that inform representation learning.

### Model architecture and training

After retrieval augmentation, EPERep employs a protein embedding pipeline *f*, which consists of ESM2^30^ and ProteinCLIP^31^, to produce initial representations for both the query and retrieved sequences:

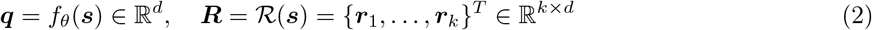

where ***q*** denotes the query protein embedding, ***R*** is the matrix of embeddings for the *k* retrieved proteins, and *d* is the dimensionality of ProteinCLIP embeddings.

To integrate contextual information from homologs, we employ a multi-head attention module^32^, which generates a fused representation of the query and retrieved proteins:

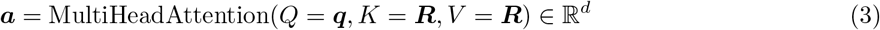

where *Q, K*, and *V* denote the query, key, and value inputs to the attention module.

To emphasize the central role of the query protein, we introduce a residual gating mechanism. Specifically, we compute a learnable scalar gate *α* that balances the fused representation ***a*** with the original query embedding ***q***:

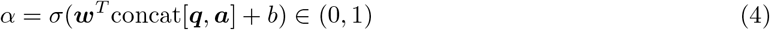

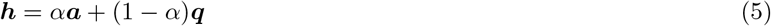

where *σ* is the sigmoid function. This gating mechanism adaptively controls the contribution of homologous context relative to the original query representation. Finally, the mixed representation ***h*** is passed through a series of fully connected layers *g*_*ϕ*_ to produce logits for each function class:

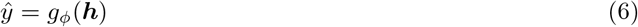

During training, we keep the ESM2 and ProteinCLIP encoders frozen and optimize all other modules with cross-entropy loss, which keeps the approach efficient and easy to scale across different function-annotation tasks. Complete hyperparameters and additional training details are provided in Supplementary Note A.3.

## Availability of data and materials

All the protein sequence and function annotation data used for model training, evaluation, and analysis were obtained from UniProt (https://www.uniprot.org/). The processed function annotation data, along with protein sequence embeddings, are available at https://www.dropbox.com/scl/fi/53fzzzmyy1upnpsjj6ei7/data.tar.gz?rlkey=dm8hs60t0j7w5bzg4nc3fshv7&st=zcoodjl5&dl=1. The source code of EPERep is available at the GitHub repository https://github.com/luo-group/EPERep.

## Funding

This work is supported in part by the National Institute of General Medical Sciences of the National Institutes of Health (R35GM150890) and a National Science Foundation (NSF) CAREER Award (No. 2442063). The authors acknowledge the computational resources provided by the CloudHub GenAI Seed Grant funded by GaTech IDEaS and Microsoft and the Delta GPU Supercomputer at NCSA of UIUC through allocation CIS230097 from the ACCESS program, which is supported by NSF grants #2138259, #2138286, #2138307, #2137603, and #2138296.

## Authors’ contributions

Y.L. conceived the ideas. S.D. and J.L. designed and performed the experiments. S.D. and J.L. analyzed the data and results. S.D. prepared the figures. S.D., J.L., and Y.L. wrote the manuscript. Y.L. supervised the project. All authors contributed ideas to the work and wrote the manuscript.

## A Supplementary Notes

### A.1 Evaluation Metrics

Considering the protein function prediction tasks are multi-label, multi-class classification problems and each sample could have multiple associated labels. Below, we describe the metrics used to evaluate the model’s performance.

#### Precision

Precision quantifies how many of the predicted positive labels are actually correct. For a single test instance, it is computed as:

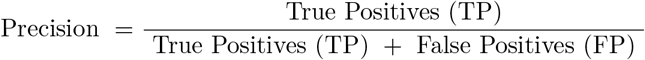

Here, TP represents the count of correctly predicted labels, and FP is the count of incorrect predictions. To calculate the overall precision across the test set, the precision for each instance is averaged: 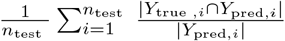, where *n*_test_ is the number of test instances, *Y*_true, *i*_ is the set of ground truth labels for the *i*-th instance, and *Y*_pred, *i*_ is the set of predicted labels for the same instance. When evaluating precision on specific subsets of labels, only the corresponding columns from the binary label matrices are considered.

#### Recall

Recall measures the model’s ability to identify all relevant labels for a given instance. For a single test sample, it is defined as 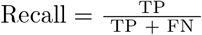. Here, FN refers to the number of relevant labels that were missed by the model. The average recall over the entire test set is calculated by summing the recall values for each test instance and dividing by the number of test samples: 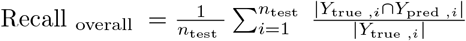

#### F1 score

The F1 score balances precision and recall, providing a harmonic mean of the two metrics. For an individual test sample, it is calculated as 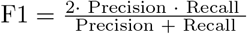. This can also be expressed in terms of TP, FP, and FN as 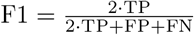. The overall F1 score for the test set is computed by averaging the F1 scores across all test samples: 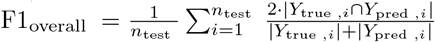

#### AUPR

The AUPR measures the area under the precision-recall curve, which reflects the trade-off between precision and recall for varying thresholds. To compute AUPR, ***Y***_prob_ and ***Y***_true_ are flattened into 1D arrays, and the precision-recall curve is calculated using precision recall curve in scikit-learn. The area under this curve is then computed using auc. For class subsets, the relevant columns of ***Y***_prob_ and ***Y***_true_ are selected, flattened, and used to calculate AUPR. These metrics ensure consistent evaluation for imbalanced multi-label classification tasks.

#### Fmax socre

The Fmax score represents the maximum F1 score achieved across a range of thresholds. For a given threshold *ρ* ∈ [0, 1], a binary prediction matrix ***Y***_pred_ is generated from the probability matrix ***Y***_prob_ by setting entries below *ρ* to 0 and the rest to 1. Using ***Y***_pred_ and the ground truth matrix ***Y***_true,_, the F1 score is calculated with micro-averaging: 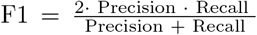. The Fmax score is defined as Fmax = max_*ρ*∈[0,1]_ F1(*ρ*). In our evaluation, thresholds are sampled at 101 equally spaced values between 0 and 1. For class subsets, the same process is applied to reduced matrices 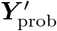 and 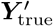.

### A.2 Baselines Methods

Aspect-Vec^3^ is a contrastive learning framework built on protein language model representations. It learns a protein sequence representation space where proteins of the same functions are pulled together, while proteins of different functions are pushed away from each other. During inference, it employs a nearest neighbor search scheme to transfer the function annotations of the most similar lookup protein to the query protein as predictions. We included Aspect-Vec for EC number, Gene3D, and Pfam predictions and used the model checkpoints provided by the paper, which are trained on the same time-split dataset of EPERep.

Protein-Vec^3^ learns multi-aspect representations via contrastive learning on the representations of Aspect-Vec. It further fuses the Aspect-Vec representations for different tasks into a unified embedding and adopts the same nearest neighbor search scheme during inference. We used Protein-Vec for all annotation tasks, including EC number, Gene3D, Pfam, and Gene Ontology.

BLAST^1^ is a common sequence search tool based on local alignment. For each annotation task, we use BLAST to search the test set against the corresponding training set, and take the function annotations of the lookup sequence with the highest bit score as predictions for each test sequence. We used BLAST for EC number and Gene3D predictions.

CLEAN^7^ is a contrastive learning framework for EC number prediction. For a fair comparison, we trained CLEAN on the same training set as our model and followed the original paper’s inference protocol (maximum separation with default hyperparameters).

HMMER^2^ is a profile-based sequence search method. We first use HHblits^5^ to build MSAs for each sequence in the lookup set against our retrieval search database, UniRef30. HMM profiles are built from the MSAs using the `hmmbuild` command. Finally, we use the `hmmsearch` command to search each query sequence in the test set against the profile database. For each task, the lookup set is the training set of EPERep.

NetGO3.0^6^ is an ensemble-based Gene Ontology (GO) prediction method. It is a combination of various function prediction methods, including GO term frequency, sequence similarity-based KNN, k-mer, and protein domain-based linear regression, protein interaction networks, and protein language model-based linear regression. It employs a learning-to-rank (LTR) model to integrate predictions from different methods as final GO term predictions.

AnnoPRO^8^ is a deep learning framework for GO prediction. It leverages a convolutional neural network to encode multi-scale protein representations and long short-term memory (LSTM)-based decoding for GO term predictions.

MSRep^4^ is a protein representation learning framework for function annotations, including EC number, Gene3D, Pfam, and Gene Ontology. MSRep learns a function-informed representation space for protein sequences where the distance between protein representations indicates their functional similarity. MSRep conducts the nearest neighbor search for function annotation transfer between query sequences and lookup sequences. MSRep is trained on the same time-based split datasets as EPERep.

### A.3 Training hyperparameters

All models of EPERep were implemented in `PyTorch Lightning` and optimized with the AdamW optimizer. Learning rates were tuned in {1 *×* 10^−5^, 5 *×* 10^−5^, 1 *×* 10^−4^} and weight decay in {0, 1 *×* 10^−2^}, with the best configuration selected on a held-out validation split via grid search. The number of retrieved sequences *k* was varied from 1 to 15, with *k* = 10 yielding the best trade-off between accuracy and efficiency. The shared embedding dimension *d* was 128, and multi-head attention was evaluated with *H* ∈ {2, 4, 8} heads. Unless otherwise stated, we report results with *k* = 10, *d* = 128, *H* = 4, and a learning rate of 1 *×* 10^−4^. Early stopping was applied based on validation AUPR with a patience of 10 epochs. All experiments were trained on NVIDIA A40 GPUs with a batch size of 1024.

## B Supplementary Figures

**Figure S1:**
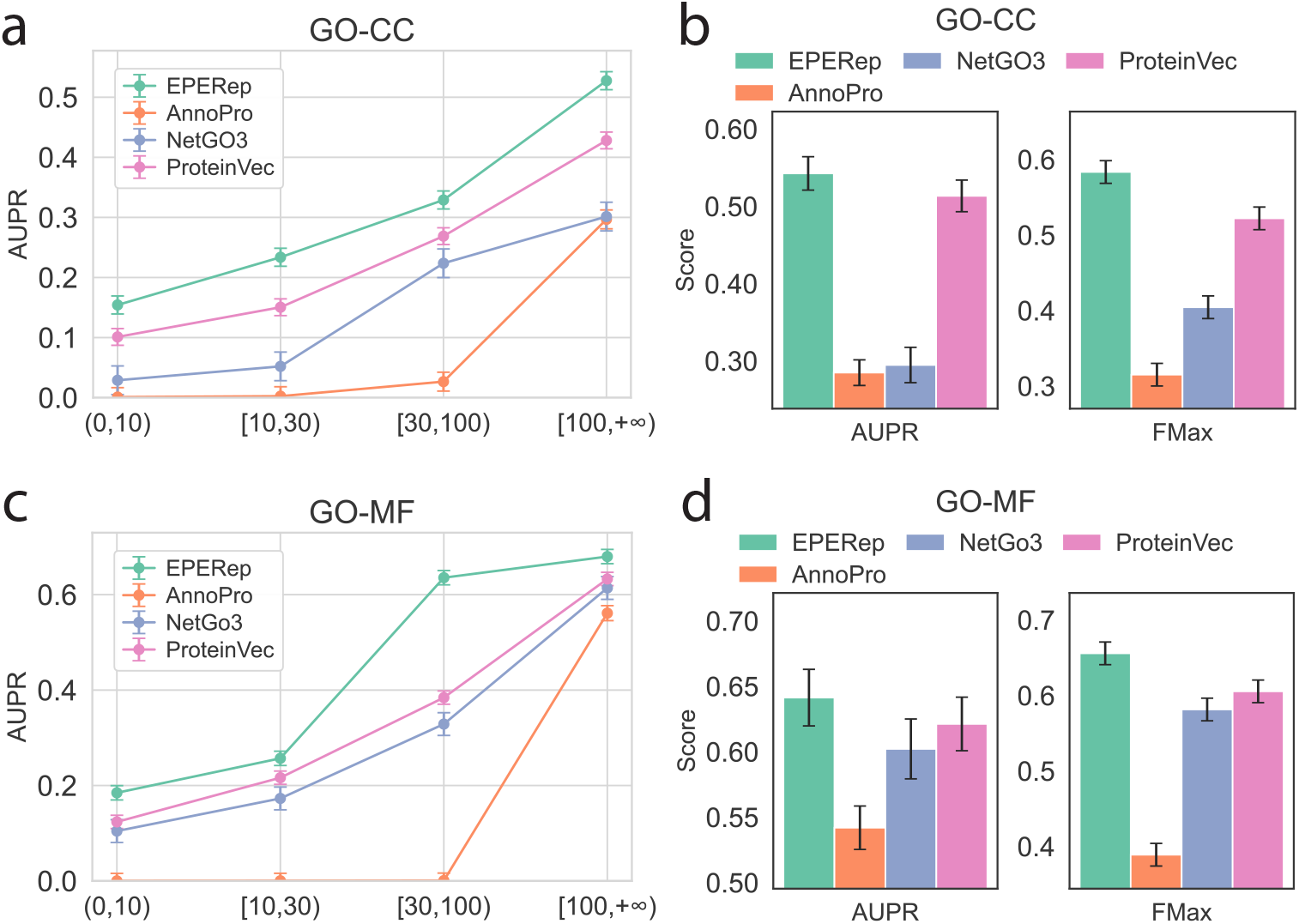
Performance of GO-BP and GO-CC. **(a)** AUPR and Fmax scores of EPERep and corresponding baselines on the GO-CC task. CC: Cellular Component. **(b)** AUPR scores of EPERep and corresponding baselines on different subsets of GO-CC task. **(c)** AUPR and Fmax scores of EPERep and corresponding baselines on the GO-MF task. MF: Molecular Function. **(d)** AUPR scores of EPERep and corresponding baselines on different subsets of GO-MF task.

## Notes

### Competing Interest Statement

The authors have declared no competing interest.

